# Detecting Heterogeneity in Population Structure Across the Genome in Admixed Populations

**DOI:** 10.1101/031831

**Authors:** Caitlin McHugh, Lisa Brown, Timothy A. Thornton

**Affiliations:** Department of Biostatistics, University of Washington, Seattle, WA 98195

**Keywords:** admixture, population structure, heterogeneity testing, local ancestry, assortative mating

## Abstract

The genetic structure of human populations is often characterized by aggregating measures of ancestry across the autosomal chromosomes. While it may be reasonable to assume that population structure patterns are similar genome-wide in relatively homogeneous populations, this assumption may not be appropriate for admixed populations, such as Hispanics and African Americans, with recent ancestry from two or more continents. Recent studies have suggested that systematic ancestry differences can arise at genomic locations in admixed populations as a result of selection and non-random mating. Here, we propose a method, which we refer to as the chromosomal ancestry differences (CAnD) test, for detecting heterogeneity in population structure across the genome. CAnD uses local ancestry inferred from SNP genotype data to identify chromosomes harboring genomic regions with ancestry contributions that are significantly different than expected. In simulation studies with real genotype data from Phase III of the HapMap Project, we demonstrate the validity and power of CAnD. We apply CAnD to the HapMap Mexican American (MXL) and African American (ASW) population samples; in this analysis the software RFMix is used to infer local ancestry at genomic regions assuming admixing from Europeans, West Africans, and Native Americans. The CAnD test provides strong evidence of heterogeneity in population structure across the genome in the MXL sample *(p = 4e −* 05), which is largely driven by elevated Native American ancestry and deficit of European ancestry on the X chromosomes. Among the ASW, all chromosomes are largely African derived and no heterogeneity in population structure is detected in this sample.

## Introduction

Technological advancements in high-throughput genotyping and sequencing technologies have allowed for unprecedented insight into the genetic structure of human populations. Population structure studies have largely focused on populations of European descent, and ancestry differences among European populations have been well studied and characterized (Novembre *et al*. 2008; Nelis *et al*. 2009). Recent studies have also investigated the genetic structure of more diverse populations, including recently admixed populations, such as African Americans (Zakharia et al. 2009; Bryc et al. 2010a) and Hispanics (Manichaikul *et al*. 2012), who have experienced admixing within the past few hundred years from two or more ancestral populations from different continents.

Both continental and fine-scale genetic structure of human populations have largely been characterized by aggregating measures of ancestry across the autosomal chromosomes. While it may be reasonable to assume that population structure patterns across the genome are similar for populations with ancestry derived from a single continent, such as populations of European descent, this may not be a reasonable assumption for recently admixed populations who have ancestries from multiple continents. For example, a previous analysis of Puerto Rican samples identified multiple chromosomal regions with large, systematic ancestry differences, as compared to what would be expected based on genome-wide ancestry, and thus providing evidence of recent selection in this admixed population (Tang *et al*. 2007). Sex-specific patterns of non-random mating at the time of or since admixture can also result in systematic differences in ancestry at genomic loci as well as across entire chromosomes, such as the X and Y chromosomes, in admixed populations. For example, a recent study compared the average ancestry on the autosomes to the X chromosome in a large sample of Hispanics and African Americans (Bryc *et al*. 2015) and highly significant differences in ancestry were detected, with increased Native American and African ancestry, respectively, on the X chromosome in the Hispanic and African American samples, and a deficit of European ancestry as compared to the autosomes.

Previous methods (Tang *et al*. 2007; Jin *et al*. 2012; Bhatia *et al*. 2014) have been proposed to identify signals of selection by detecting genomic regions in admixed populations that exhibit unusually large deviations in ancestry proportions compared to what is expected based on genome-wide ancestry. For assessing significance, however, these methods require strong assumptions about the evolution of the admixed population of interest, which will generally be partially or completely unknown, including (1) the relative contribution from each of the ancestral populations to the gene pool at the time of the admixture events, (2) the number of generations since the admixture events, (3) an assumed effective population size, and (4) random mating. Significance is then assessed either analytically or through simulation studies based on these evolutionary assumptions. Mis-specification of these assumptions, however, can result in false positives due to an incorrect null distribution, and regions of the genome that appear to have large ancestry differences are actually not significantly different from what would be expected when sampling variation, genetic drift after admixture, and potential bias in local ancestry estimation is appropriately taken into account (Bhatia *et al*. 2014).

Here, we consider the problem of detecting heterogeneity in ancestry across the genome in admixed populations. We propose the Chromosomal Ancestry Differences (CAnD) test for the identification of chromosomes that harbor genomic regions with significantly different proportional ancestry as compared to the rest of the genome. For each sampled individual, CAnD incorporates ancestry inferred at genomic regions using local ancestry methods, such as HAPMIX (Price *et al*. 2009) or RFMix (Maples *et al*. 2013), and tests for systematic differences in genetic contributions to the chromosomes from the underlying ancestral populations. The CAnD method takes into account correlated ancestries among chromosomes within individuals for improved power, and the method can be used for the detection of ancestry differences among the autosomes, as well as between the autosomes and the X chromosome.

We perform simulation studies using real genotype data from Phase III of the HapMap Project (Altshuler *et al*. 2010) to evaluate the type I error rate and power of CAnD. We also apply CAnD to the HapMap Mexican Americans from Los Angeles, California (MXL) and African Americans from Southwest U.S.A. (ASW) population samples for the detection of heterogeneity in population structure. In this analysis, RFMix is used to infer European, Native American, and African ancestry at genomic locations across the autosomes and the X chromosome using RFMix. In both simulation studies and in HapMap, we compare heterogeneity testing of ancestry for the autosomes and the X chromosome with CAnD to the t-test that does not account for ancestry correlations among chromosomes of an admixed individual.

## Methods

### Chromosomal and Genome-wide Ancestry Measures

Let *n* be the number of unrelated individuals sampled from a population derived from *K* ancestral subpopulations. For individual *i*, *i* ∈ {1,…, *n*}, we define the overall, or genome-wide, ancestry of *i* as measured across the autosomal and X chromosomes. (For males, ancestry on the Y chromosome could also be included when calculating genome-wide ancestry if this information is available). Quantitatively, we denote the genome-wide ancestry vector for individual *i* as **a***_i_* = (*a_i_*_1_,…,*a_iK_*)*^T^*, where is the proportion of ancestry from subpopulation *k* for individual *i, a_ik_*) ≥ 0 for all *k*, and 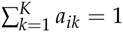.

Consider the set 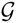 of autosomal and X chromosomes, i.e., 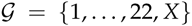. Denote the genetic ancestry for individual *i* on a particular chromosome 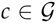 as 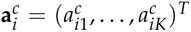. For each chromosome *c*, denote 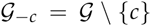 to be the set of all chromosomes excluding *c*, i.e., 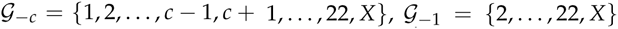 and 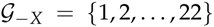. Define 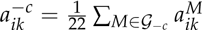 to be the mean of all chromosomal ancestries with chromosome *c* excluded for subpopulation *k* and individual *i*. Note for individual *i*, 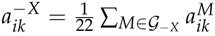 is the average autosomal ancestry for subpopulation *k*. We define 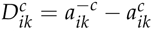 to be the difference in ancestry between a given chromosome *c* and the mean ancestry of all other other chromosomes in individual *i* for subpopulation *k*. We denote 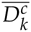 to be the mean of the 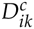 values across all individuals *i* ∈ {1,…, *n*}.

### The CAnD Test

Consider the previously defined set 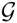 consisting of the autosomal and X chromosomes. To test for heterogeneity in ancestry from subpopulation *k* among a subset 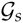 of 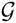, where 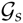 could also be 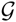 i.e., 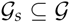, that contains *m* chromosomes, we first calculate a statistic 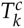 for each chromosome 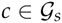 that is the mean of the standardized proportional ancestry differences for population k between *c* and the pooled average ancestry of all other chromosomes in 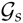 within each of the *n* sampled individuals, where

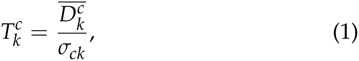

and *σ_ck_* is the standard deviation of 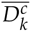 (defined in the previous subsection). Under the null hypothesis of no ancestry differences among the *m* chromosomes, 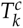 approximately follows a normal distribution with mean 0 and variance 1 for each 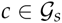, and the multivariate statistic

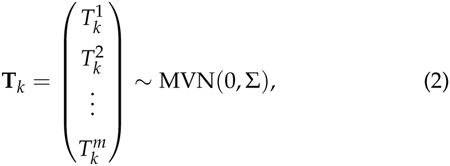

where Σ is the *m × m* covariance matrix of *T_k_*, allowing for correlation among the 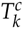 statistics. To test for heterogeneity in ancestry from population *k* among the *m* chromosomes in 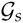, we propose the chromosomal ancestry differences (CAnD) test statistic

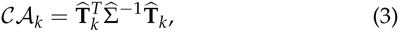

where 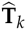 is T*_k_* calculated with 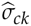 evaluated at *σ_ck_* for each chromosome *c*, and 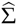 is an estimate of Σ. Under the null hypothesis, 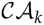 approximately follows a χ^2^ distribution with *m* degrees of freedom. Details about the estimators 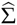 and 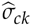 for Σ and *σ_ck_*, respectively, that we propose are given in Appendix A.

### Simulation Studies

In order to assess type I error and power of the CAnD method, we performed simulation studies using real data from the HapMap CEU (Utah residents with ancestry from northern and western Europe from the Centre d’Étude du Polymorphisme Humain collection) and YRI (Yoruba in Ibadan, Nigeria) populations. Each simulated replicate consisted of simulated chromosomes for 50 admixed individuals that were derived from 118 CEU and YRI haplotypes on chromosomes 1 and 2, where the chromosomal haplotypes consisted of 5,000 evenly spaced markers (Altshuler *et al*. 2010) across the chromosome.

Each simulated admixed individual *i ∈* {1,…,50} has admixture vectors for chromosomes 1 and 2 of the form 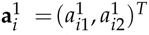 and 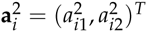, respectively, where 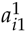 and 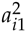 are the population 1 ancestry proportions on chromosomes 1 and 2, respectively, and 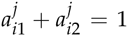 for *j =* 1,2. We denote CEU and YRI to be populations 1 and 2, respectively, in the simulation study, and proportional CEU ancestry on chromosome 1 for individual *i* is 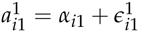 where 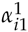 is drawn from uniform distribution on [0.05,0.45] and 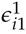 is drawn from a *N*(0, 8.2*e* − 04) distribution. The variance of 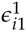 corresponds to an estimate of the average variance across the autosomal chromosomes for European ancestry within admixed individuals from the HapMap MXL. For chromosome 2, 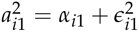, where 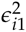 is a random ancestry effect for chromosome 2 that follows a *N*(*μ*, 8.2*e −* 04) distribution, where 0 ≤ |*μ*| ≤ 1. Under the null hypothesis, *μ =* 0, i.e., there is no difference in mean ancestry between chromosomes 1 and 2, and |*μ*| > 0 under the alternative hypothesis. Each chromosome 1 for individual *i* is constructed from the CEU and YRI haplotypes, where the chromosome has proportions 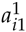 and 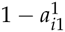, respectively, from a randomly drawn CEU haplotype and a randomly drawn YRI haplotype. The two copies of chromosome 2 for individual *i* are similarly obtained.

For each simulated individual, chromosomal-wide ancestry proportions were estimated from the genotype data using the FRAPPE software program (Tang *et al*. 2005), which uses a likelihood-based model to infer each individual’s ancestry proportions. Included as reference samples in the FRAPPE runs were 58 CEU and 57 YRI HapMap samples, and the number of reference populations was set to two. The reference samples used for the FRAPPE analyses were different from those used to simulate the admixed individuals’ genotypes. With the resulting FRAPPE proportions, we implemented the CAnD method to identify heterogeneity in population structure across two chromosomes. A variety of *μ* values were considered for the assessment of type I error and power at different significance levels.

### HapMap MXL and ASW

We considered detection of heterogeneity in ancestry across the genome in unrelated HapMap MXL and ASW samples. REAP (Thornton *et al*. 2012) was used to infer both known and cryptic relatedness in the MXL and ASW, and a subset of 53 MXL individuals and a subset of 45 ASW individuals with inferred relationships less than third degree were identified and included for the ancestry heterogeneity analysis. Of the unrelated subset of 53 MXL individuals, there were eight singletons, 20 families with two individuals included and one family with three individuals. Among the 45 unrelated ASW individuals, there were 23 singletons and 11 families with two individuals that were included. There were 27 females and 26 males in the unrelated HapMap MXL subset, and 25 females and 20 males in the unrelated HapMap ASW subset. We also performed CAnD tests stratified by sex to determine if there was any bias in the results due to copy number differences in the X chromosome for males and females.

We used the RFMix software (Maples *et al*. 2013) to estimate local ancestry across the autosomes and the X for all HapMap MXL and ASW samples. RFMix allows for more than two ancestral subpopulations and in both the HapMap MXL and ASW analyses, and we assumed ancestral contributions from African, European and Native American populations. The HapMap CEU and YRI samples were included as the reference population panels in the local ancestry analysis for European and African ancestry, respectively, and the Human Genome Diversity Project (HGDP) (Li *et al*. 2008) samples from the Americas were included as the reference population panel for Native American ancestry. All samples were phased and sporadic missing genotypes were imputed using the BEAGLE v.3 software (Browning and Browning 2007). Recombination maps for each chromosome were downloaded from the HapMap website (Altshuler *et al*. 2010) and were converted to Human Genome Build 36. There was no phasing conducted on the X for males since a male only has one X chromosome. Only SNPs that were genotyped in both the HapMap and HGDP datasets were considered in the local ancestry analysis. For local ancestry on the X chromosome, SNPs on the non-pseudoautosomal regions, where there is no homology between the X and Y chromosomes, were considered.

We compared CAnD when using global ancestry for each chromosome estimated using the FRAPPE software (Tang *et al*. 2005) to CAnD when using local ancestry estimated across the chromosomes with RFMix. For each chromosome, a supervised global ancestry analysis was conducted separately for the HapMap MXL and ASW population samples with FRAPPE. The number of ancestral populations was set to three and the same reference population samples used in the RFMix local ancestry analysis were also used with FRAPPE. Since males only have one allele at each of the X chromosome SNPs, one of the alleles at an X-linked SNP was coded to be missing in the FRAPPE analysis, although we found that coding male genotypes as homozygous in the FRAPPE analysis yielded nearly identical results.

## Results

### Assessment of Type I Error

In the simulation studies for detecting ancestry heterogeneity, FRAPPE was first used to estimate proportional ancestry on chromosomes 1 and 2 for each simulated admixed individual. To ensure that the FRAPPE estimates were accurate when using unphased genotypes from 5,000 SNPs on a chromosome, we first compared the FRAPPE ancestry estimates to the simulated ancestry. The differences between the FRAPPE estimates and the simulated ancestry proportion values have mean of −5.147e-06 (SD=0.018), indicating FRAPPE can accurately estimates chromosomal ancestry proportions when using a set of 5,000 markers (Figure S1).

To assess the type I error rate of CAnD, we simulated admixed chromosomes for 50 sampled individuals under the null hypothesis of no ancestry differences among the chromosomes, on average. The empirical type I error rates for the CAnD test at the *α =* 0.01, 0.005, and 0.001 significance levels calculated using 5,000 simulated replicates are given in Table 1. The CAnD test is properly calibrated for all significance levels considered. Empirical type I error rates are not significantly different from the nominal levels, as can be seen from the 95% confidence intervals given in the table.

**Table 1.**
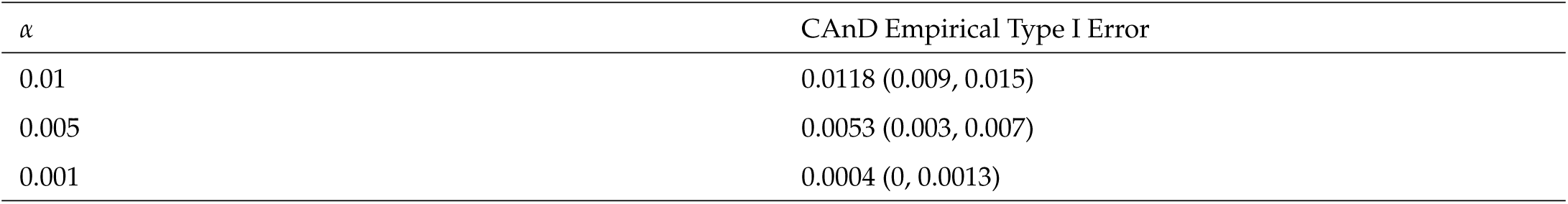
Empirical Type I Error. CAnD Empirical Type I Error (95% CI) at significance levels *α =* 0.01, 0.005, and 0.001 based on 5,000 simulated replicates. This simulation setting was conducted under the null hypothesis where the randomly drawn ancestry proportions of an admixed individual are the same for both chromosomes 1 and 2.

### Power Evaluation and Comparison

We evaluated the power of the CAnD method for an admixed sample of 50 individuals. The values of *μ*, the mean difference in ancestry between chromosomes 1 and chromosome 2, ranged from 0.005 and 0.25. We also compared the power of CAnD to a pooled t-test that ignores the correlation of ancestry across chromosomes within an individual. Although ancestry across chromosomes are not independent within an individual, we present this method for comparison to CAnD as it has been used in previous studies for the testing of ancestry differences between the autosomal chromosomes and the X chromosome in admixed populations (Bryc *et al*. 2015).

Empirical power results at the *α =* 0.01 significance level using the CAnD and pooled t-test are given in Figure 1. CAnD has higher power than the pooled t-test for all values of *μ* considered, and significantly higher values for low to moderate values of *μ*. For example, there is essentially no power to detect a mean difference in ancestry of 5% between the two chromosomes with the pooled t-test, while CAnD has power that is close to 1. The loss in power with the pooled t-test is due to the test not accounting for the correlation in ancestry between chromosomes within an individual. We recommend the CAnD test over the pooled t-test for improved power to detect ancestry differences among chromosomes.

**Figure 1.**
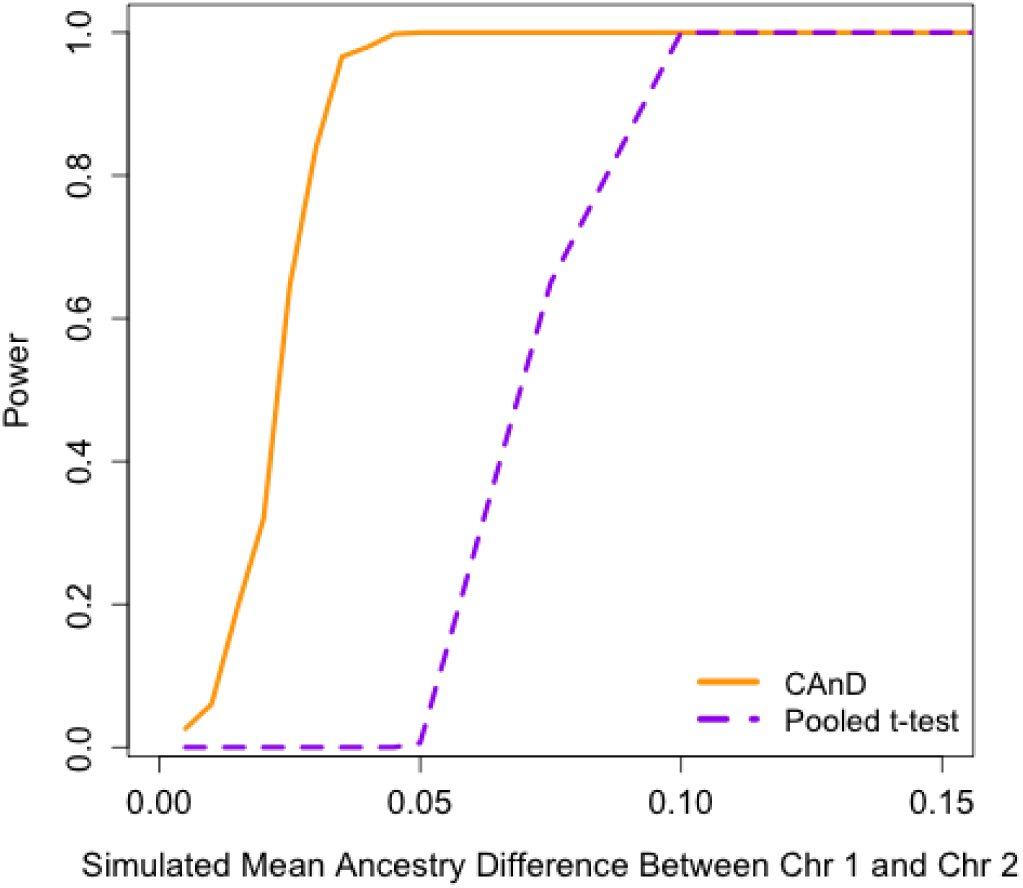
Power of the CAnD Test in Simulated Data. The proportion of tests rejected at a significance level of 0.01 when using the CAnD method as compared to the pooled t-test under increasing differences in ancestry proportion between chromosomes. For each simulated ancestry proportion difference, the proportion of tests rejected was calculated from 500 independent simulations of 50 samples each.

### HapMap ASW Ancestry

The predominant genome-wide ancestry in all 87 HapMap ASW subjects is African. Table 2 shows the mean and SD of the local ancestry estimates by chromosome in each of the ancestral populations and accompanying Figure 2A shows violin plots of the local ancestry results by chromosome. RFMix estimated 11 individuals to have no European ancestry on the X chromosome, and the maximum European ancestry on the X chromosome is 0.67. On the other hand, nine individuals are estimated to have an X chromosome entirely of African ancestry, where the proportion ranges from 0.33 to 1. We see these patterns in the barplots shown in Figure 3A which displays the proportion ancestry for each sample. Across both the autosomes and the X chromosome, the proportion Native American is quite small over all samples. Fifty-seven individuals are estimated to have no Native American ancestry on the X chromosome. While the proportion of Native American ancestry is larger on the autosomes than the X chromosome, on average, it remains small in magnitude and we conclude that Native American ancestry is negligible in this sample of individuals. Furthermore, we detect more African and less European ancestry on the X chromosome than the autosomes, overall.

**Table 2.** Summary of Local Ancestry Estimates by Chromosome. Mean (SD) of local ancestry estimates by chromosome, stratified by the ASW and MXL HapMap population samples.

| Chr | ASW |  |  | MXL |  |  |
| --- | --- | --- | --- | --- | --- | --- |
|  | African | European | Native American | African | European | Native American |
| X | 0.82 (0.139) | 0.163 (0.136) | 0.017 (0.047) | 0.0396 (0.0521) | 0.387 (0.245) | 0.574 (0.248) |
| Autosomal-Wide | 0.783 (0.0861) | 0.202 (0.0808) | 0.0150 (0.0382) | 0.0489 (0.0182) | 0.508 (0.149) | 0.444 (0.148) |
| 1 | 0.762 (0.13) | 0.228 (0.131) | 0.00962 (0.0354) | 0.047 (0.0389) | 0.525 (0.192) | 0.428 (0.191) |
| 2 | 0.789 (0.132) | 0.201 (0.128) | 0.0106 (0.0241) | 0.0457 (0.0379) | 0.514 (0.195) | 0.44 (0.188) |
| 3 | 0.769 (0.155) | 0.221 (0.154) | 0.0102 (0.0369) | 0.0462 (0.0345) | 0.514 (0.18) | 0.439 (0.183) |
| 4 | 0.807 (0.136) | 0.177 (0.134) | 0.0164 (0.0398) | 0.0461 (0.0408) | 0.47 (0.212) | 0.484 (0.206) |
| 5 | 0.786 (0.149) | 0.199 (0.146) | 0.0148 (0.0536) | 0.0539 (0.05) | 0.528 (0.2) | 0.418 (0.188) |
| 6 | 0.774 (0.167) | 0.201 (0.15) | 0.0257 (0.0696) | 0.0555 (0.0502) | 0.5 (0.179) | 0.445 (0.177) |
| 7 | 0.804 (0.125) | 0.184 (0.117) | 0.012 (0.0539) | 0.056 (0.047) | 0.524 (0.193) | 0.42 (0.188) |
| 8 | 0.785 (0.163) | 0.201 (0.16) | 0.0141 (0.0419) | 0.0397 (0.0349) | 0.504 (0.187) | 0.456 (0.179) |
| 9 | 0.772 (0.12) | 0.21 (0.116) | 0.0175 (0.0506) | 0.0499 (0.0521) | 0.489 (0.21) | 0.462 (0.213) |
| 10 | 0.785 (0.145) | 0.205 (0.14) | 0.00997 (0.047) | 0.059 (0.066) | 0.502 (0.189) | 0.439 (0.183) |
| 11 | 0.778 (0.141) | 0.212 (0.139) | 0.00953 (0.0255) | 0.0402 (0.0422) | 0.525 (0.202) | 0.435 (0.201) |
| 12 | 0.779 (0.142) | 0.202 (0.137) | 0.0186 (0.0625) | 0.0501 (0.0488) | 0.511 (0.18) | 0.439 (0.177) |
| 13 | 0.804 (0.152) | 0.18 (0.149) | 0.0165 (0.032) | 0.0488 (0.0424) | 0.523 (0.199) | 0.428 (0.201) |
| 14 | 0.802 (0.162) | 0.183 (0.155) | 0.015 (0.0577) | 0.0559 (0.0643) | 0.47 (0.217) | 0.474 (0.219) |
| 15 | 0.817 (0.141) | 0.172 (0.138) | 0.0109 (0.0399) | 0.0382 (0.0452) | 0.528 (0.182) | 0.434 (0.179) |
| 16 | 0.778 (0.192) | 0.201 (0.183) | 0.0207 (0.0631) | 0.0456 (0.0449) | 0.498 (0.205) | 0.457 (0.209) |
| 17 | 0.772 (0.156) | 0.207 (0.145) | 0.0208 (0.079) | 0.041 (0.043) | 0.533 (0.195) | 0.426 (0.192) |
| 18 | 0.772 (0.21) | 0.21 (0.196) | 0.0184 (0.0605) | 0.0501 (0.047) | 0.537 (0.207) | 0.413 (0.198) |
| 19 | 0.78 (0.154) | 0.213 (0.155) | 0.00745 (0.0189) | 0.0654 (0.0809) | 0.506 (0.208) | 0.429 (0.202) |
| 20 | 0.801 (0.167) | 0.187 (0.152) | 0.0125 (0.0611) | 0.0541 (0.0616) | 0.52 (0.194) | 0.426 (0.195) |
| 21 | 0.76 (0.191) | 0.22 (0.186) | 0.0202 (0.0748) | 0.0451 (0.0529) | 0.475 (0.235) | 0.48 (0.232) |
| 22 | 0.747 (0.209) | 0.234 (0.206) | 0.0188 (0.0671) | 0.0419 (0.0506) | 0.47 (0.196) | 0.488 (0.204) |

**Figure 2.**
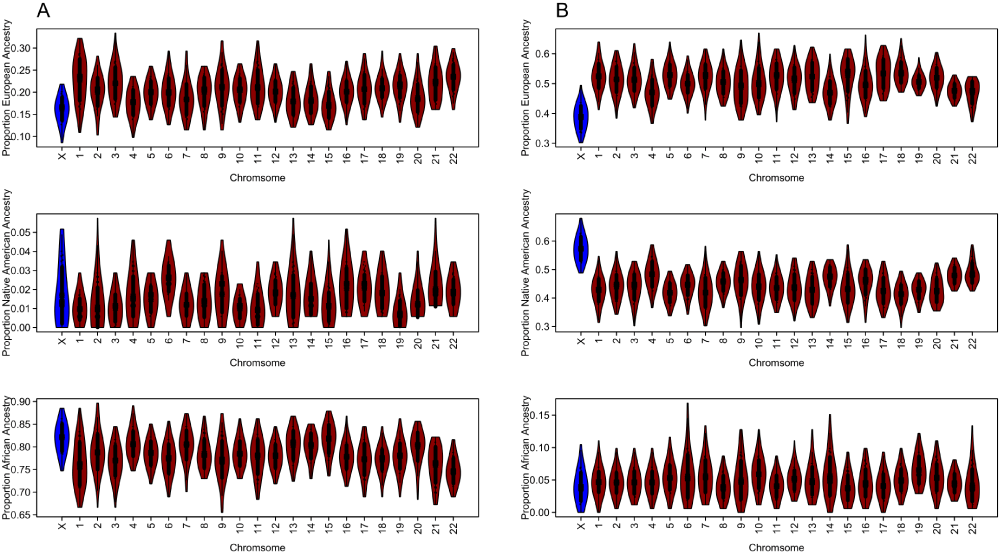
Local Ancestry Estimates by Chromosome. Chromosomal averaged local ancestry estimates for HapMap individuals using the RFMix software. Ancestry was estimated for each marker then averaged across chromosomes. (A): Estimates for 87 HapMap ASW individuals. (B): Estimates for 86 HapMap MXL individuals. The reference samples for the European and African ancestries were HapMap CEU and YRI individuals, while the HGDP samples from the Americas were references for the Native American ancestry.

**Figure 3.**
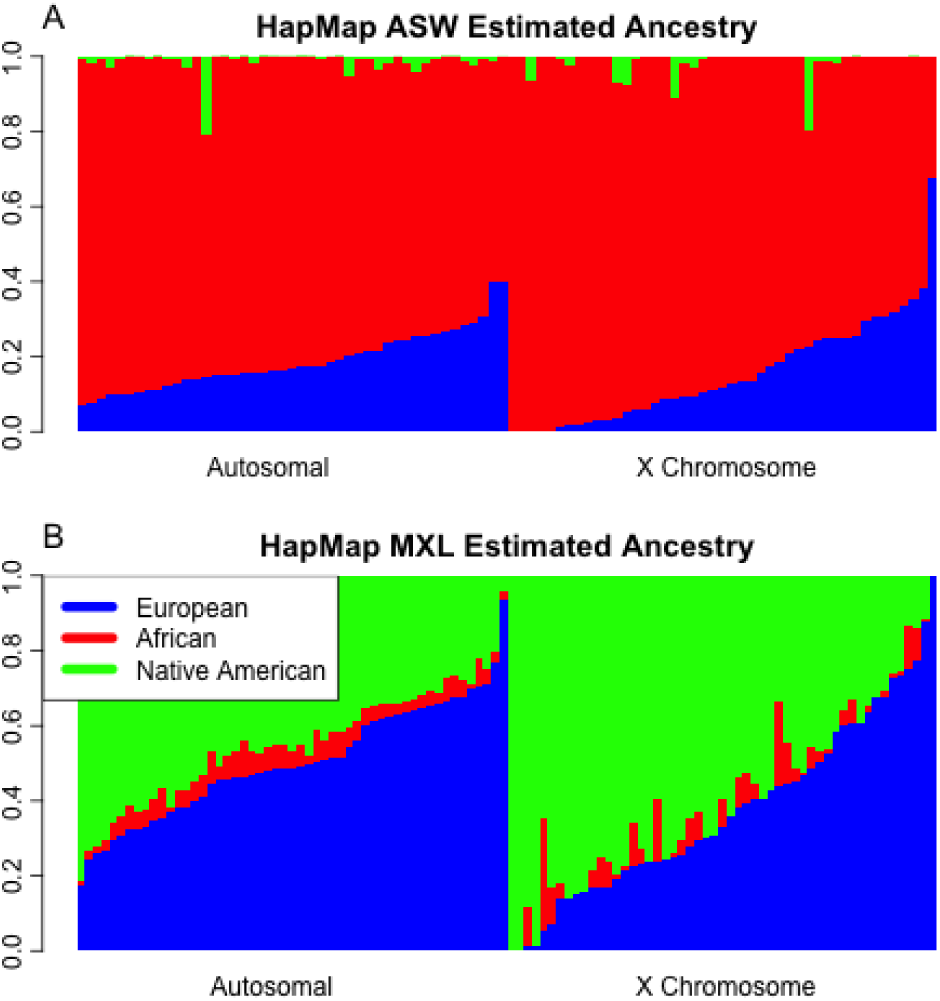
Barplots of RFMix Results. Local ancestry estimates for HapMap individuals using the RFMix software. Each individual is represented by a vertical bar, where the European, African and Native American ancestries are colored with blue, red, and green, respectively. The two panels represent the autosomal and X chromosome average. (A): Estimates for 87 HapMap ASW individuals. (B): Estimates for 86 HapMap MXL individuals. The reference samples for the European and African ancestries were HapMap CEU and YRI individuals, while the HGDP samples from the Americas were references for the Native American ancestry.

We calculated the correlation of ancestry proportions across the autosomes and X chromosome for each ancestral subpopulation. Correlation between the autosomal and X chromosome Native American ancestry is highest at 0.78. The European and African correlations between autosomal and X chromosome proportions are 0.20 and 0.17, respectively.

### HapMap MXL Ancestry

From our local ancestry analysis of the 86 HapMap MXL individuals, we found the predominant ancestries to be European and Native American, as expected based on previously reported results (Bryc *et al*. 2015; Thornton *et al*. 2012), with African ancestry being quite modest with little variation. Table 2 shows the mean and SD of the average local ancestry estimates by chromosome and averaged across the autosomes within the MXL samples. Interestingly, Native American ancestry is highest on the X chromosomes, with a mean of 57.4% (SD=24.8%), while for the autosomes, European ancestry is highest with a mean of 50.8% (SD=14.9%). African ancestry on the autosomes and the X chromosome, however, are quite similar, with mean values of 4% and 5%, respectively. Figure 2B shows violin plots by chromosome of the RFMix local ancestry estimates in the MXL samples. The plots illustrate the marked increase in proportional European ancestry across the autosomes, and, correspondingly, a decrease in proportion Native American ancestry on the autosomes as compared to the X chromosome. Estimates of ancestry on chromosome 21 and 22 are less variable than estimates across other chromosomes. Figure 3B shows barplots of the ancestral proportions within each individual. The proportion of both European and Native American ancestries on the X chromosome ranges from 0 to 1. The range and variation of the European and Native American ancestries on the X chromosome are larger than those estimated across the autosomes. Furthermore, Native American and European ancestries on the X chromosome are almost perfectly negatively correlated (corr=−0.98).

We also calculated correlation in autosomal and X chromosome ancestries. The correlation between the autosomal and X chromosome European ancestry is 0.71 and is the highest, and the Native American correlation is 0.67. With a correlation of 0.03, there is essentially no African ancestry correlation between the autosomes and the X chromosome, which likely is attributed to the small contribution of African ancestry to the HapMap MXL.

There is one male MXL individual who has an X chromosome that inferred to be completely Native American derived. The phased RFMix results of this individual’s mother indicates that one of her X chromosomes is entirely Native American while her other X chromosome is 69% Native American and 31% European, with five ancestry switches on the chromosome.

### Ancestry Heterogeneity Testing in HapMap MXL and ASW

Figure 4 shows histograms of the mean difference between the autosomal and X chromosome ancestry proportions for the sets of 45 unrelated ASW (Figure 4A) and 53 unrelated MXL (Figure 4B) individuals, with a smoothed density line overlaid. The mean difference in European ancestry between the autosomes and the X chromosome is 0.12, and the mean difference for Native American ancestry is −0.13. Based on our simulation studies, we expect to have high power to detect such large differences in ancestry between the autosomes and the X chromosome for a sample of this size. For the ASW samples, however, the mean difference between the X chromosome and the autosomes for the two predominant ancestries, African and European, is 0.04, which is much smaller than the predominant ancestry differences observed in the MXL. We expect the power to detect a mean difference in ancestry between the X and the autosomes in the ASW to be much lower, as compared to the MXL, due to the both smaller mean ancestry differences and smaller sample size.

**Figure 4.**
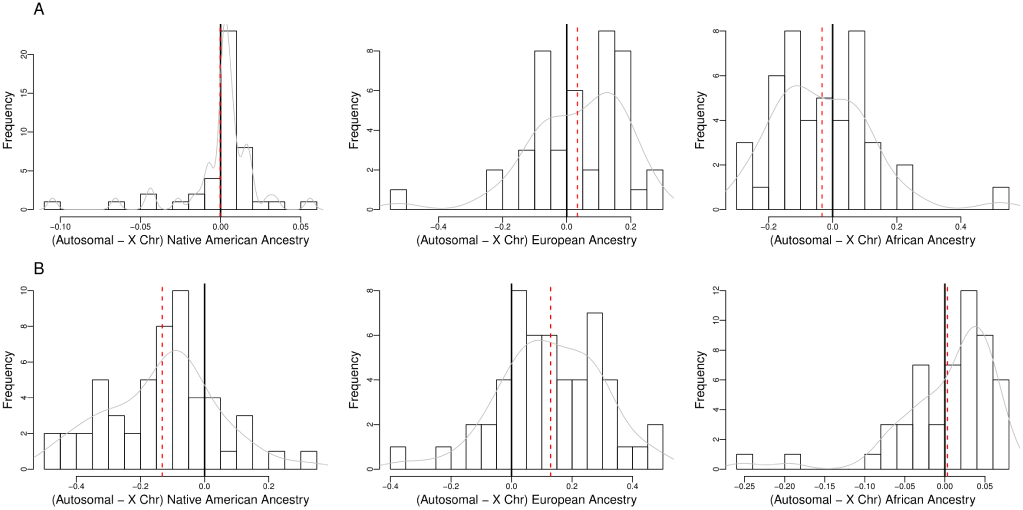
Difference in Autosomal and X Chromosome Ancestry, by Subpopulation. Histograms of the difference in autosomal and X chromosome ancestry proportions among the (A): 45 unrelated HapMap ASW and (B): 53 unrelated HapMap MXL samples. The dashed line indicates the mean difference, whereas the solid line indicates zero. A smoothed density line is overlaid on each histogram.

We applied the CAnD test to a set of 53 unrelated MXL samples. The genome-wide combined CAnD p-values are 0.592, 4.01e-05 and 9.57e-06 for the African, European and Native American ancestries, respectively. To understand which chromosomes are driving the significance found in the European and Native American ancestries, Figure 5 shows, by chromosome, the unadjusted (Figure 5A) and Bonferroni-adjusted (Figure 5B) p-values from the CAnD test in the HapMap MXL for the three ancestries. Chromosome 7 and the X chromosome have a larger proportion of Native American ancestry as compared to the mean Native American ancestry of all other chromosomes pooled together, before adjustment for multiple testing. The same result holds for the X chromosome when considering European ancestry. Chromosome 8 has a larger proportion of African ancestry than a pool of all other chromosomes. After the Bonferroni multiple testing correction, the X chromosome remains significant in the European and Native American ancestries. No other chromosomes obtain statistical significance after correction for multiple testing. Ancestry as estimated from the X chromosome is statistically significantly different from the ancestry estimates across any and all of the autosomes.

**Figure 5.**
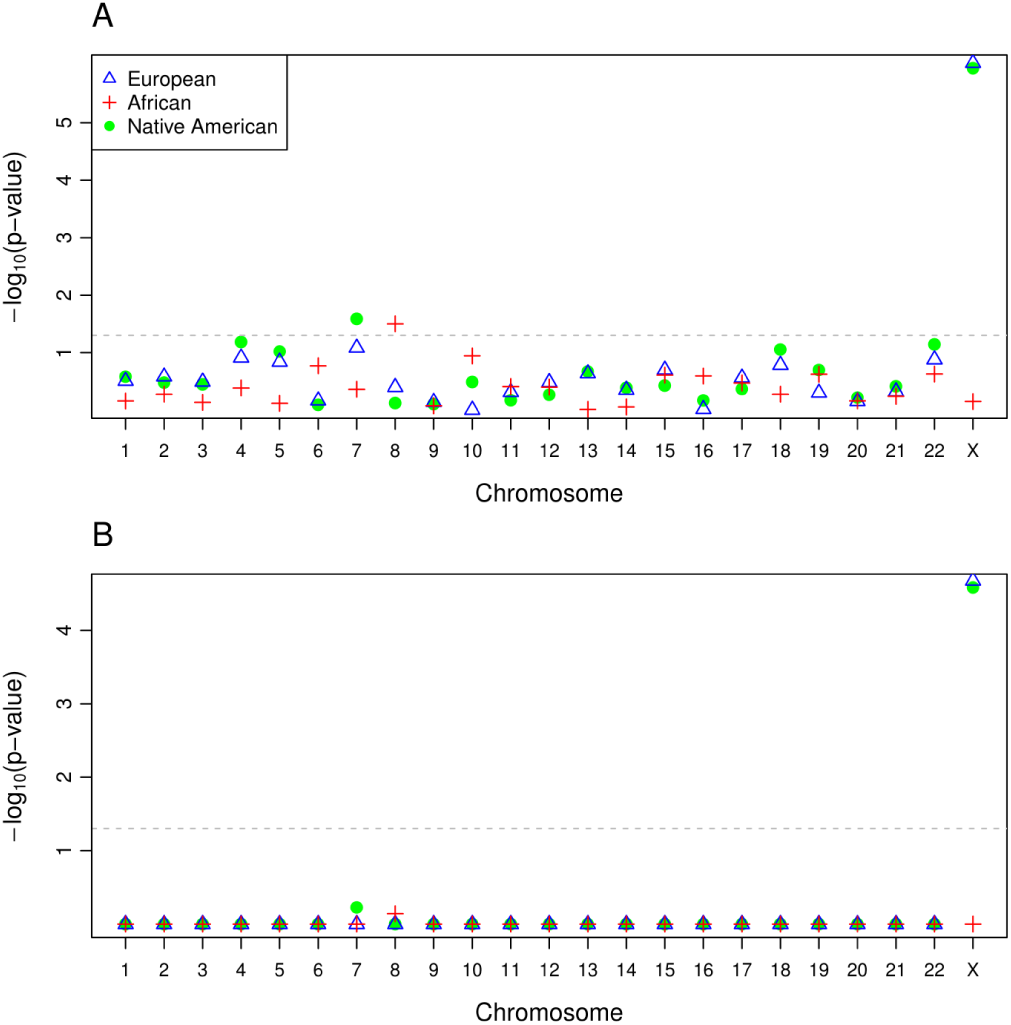
Unadjusted and Adjusted P-values from the CAnD Test in the HapMap MXL Samples. (A): Unadjusted and (B): adjusted p-values by chromosome obtained from the CAnD test comparing the estimated ancestry for each chromosome with the mean ancestry of all remaining chromosomes, including the X chromosome, for the African, European and Native American ancestries in the HapMap MXL samples. The adjusted p-values were calculated using the Bonferroni multiple testing correction.

CAnD applied to the set of 45 unrelated ASW samples yielded no significant results with genome-wide combined p-values of 0.122, 0.0858, 0.243 for the African, European and Native American ancestries, respectively (Figure S2). As previously mentioned, the autosomes and the X chromosome are predominantly African derived in the ASW, and a larger sample size is needed to achieve enough power to detect the smaller ancestry differences among chromosomes in the ASW. Indeed, in much larger population-based samples of African Americans (Bryc *et al*. 2015, 2010a), increased African ancestry and decreased European ancestry has been reported for the X chromosome as compared to the autosomes.

To assess whether inclusion of the X chromosome biased the CAnD results for the autosomal chromosomes within the HapMap MXL individuals, we performed the analysis using only the autosomes. When excluding the X chromosome, African ancestry on chromosome 8 remains significant and Native American ancestry on chromosomes 4 and 22 are significant at a 0.05 threshold (Figure S3), similar to the results when the X chromosome is included in the CAnD analysis. After correction for multiple testing using the Bonferroni procedure, however, no estimates remain significant, indicating that the significance in heterogeneity detected with CAnD is being driven by ancestry differences between the X chromosome as compared to the autosomes.

Previous studies have identified a significant difference between autosomal and X chromosome ancestry proportions in individuals from admixed populations (Bryc *et al*. 2015), where this difference has been assessed using a pooled t-test that ignores the correlation of ancestries among the autosomes and the X chromosome within an individual. We compare the performance of the pooled t-test to the CAnD test for detecting differences in ancestry between the X chromosome and the autosomes in the HapMap MXL samples. The pooled t-test finds significant differences in European ancestry and Native American ancestry between the autosomes and the X chromosome, with a p-value of 0.001 for both analyses. In comparison, the CAnD p-values comparing mean European ancestry and Native American ancestry on the X chromosome are 9.17e-07 and 1.13e-06, respectively, which is more than three orders of magnitude smaller than the pooled t-test. No significant differences in African ancestry were found using either method.

### Comparison of CAnD Results Using Local Versus Global Ancestry Estimates

We performed a CAnD analysis in the HapMap MXL and ASW using ancestry estimates for each chromosome with FRAPPE that uses unphased genotype data and assumes independent markers. We compare these results to the CAnD results reported in the previous subsection that used local ancestry estimates from RFMix, which takes into account LD among SNPs and requires phased genotype data. With the FRAPPE estimates for the ASW, no chromosomal ancestry differences were detected with CAnD, similar to the CAnD analysis results with local ancestry estimates from RFMix. Interestingly, we found that the CAnD results are slightly more significant for the MXL when using ancestry estimates from FRAPPE as compared to the estimates from RFMix, particularly for detecting differences in European ancestry across the genome (Figure S4). Inference on population structure heterogeneity in the HapMap ASW and MXL, however, is qualitatively the same with CAnD when using either local ancestry versus global ancestry estimates from RFMix or FRAPPE, respectively.

We also compared autosomal-wide and X chromosome ancestry estimates from RFMix and FRAPPE using genotype data for the HapMap MXL and ASW population samples. Table 3 shows the correlation of the ancestry estimates from the methods for each ancestral subpopulation. For the two predominant ancestries in the MXL (European and Native American) and ASW (African and European), the correlation between the ancestry estimates for the autosomes from RFMix and FRAPPE are all greater than 0.99, and is 0.95 or greater for the X chromosome. As previously mentioned, there is very little Native American ancestry and African ancestry in the ASW and MXL, respectively. Nevertheless, with a correlation of 0.99, Native American ancestry estimates on the autosomes are nearly perfectly correlated between RFMix and FRAPPE, and the correlation between the estimates is 0.90 for Native American ancestry on the X chromosome in the ASW. For proportional African ancestry in the MXL, the correlation between the two estimates is 0.893 for the autosomes and 0.93 for the X chromosome. So, for the predominant ancestries in the MXL and ASW, there appears to be little difference in estimating autosomal ancestries with FRAPPE or by averaging local ancestry estimates from RFMix. There is high concordance between the methods for the predominant ancestry in ASW and MXL for the X chromosome as well. In general, there is less concordance between the methods when estimating proportional ancestries from populations with relatively small contributions to the admixed population, and local ancestry estimates, such as RFMiX, are likely more accurate in inferring low levels of ancestral contribution, than global ancestry methods, such as FRAPPE.

**Table 3.** Correlation of Ancestry Estimates. Correlation between ancestry estimates from RFMix and FRAPPE, stratified by autosomal and X chromosome estimates, in each of the population samples.

|  | Autosomal |  | X Chromosome |  |
| --- | --- | --- | --- | --- |
|  | ASW | MXL | ASW | MXL |
| African | 0.9990 | 0.8932 | 0.9697 | 0.9256 |
| European | 0.9979 | 0.9935 | 0.9548 | 0.9878 |
| Native American | 0.9963 | 0.9940 | 0.9001 | 0.9898 |

### Assortative Mating for Ancestry in the HapMap MXL

The CAnD test identified significant heterogeneity in ancestry among the HapMap MXL chromosomes. Systematic differences in ancestry at genomic loci on chromosomes can be due to sex-specific patterns of non-random mating at the time of or since admixture. We investigated assortative mating between pairs of individuals in the HapMap MXL for which there is a documented offspring; there are 24 such pairs. However, we excluded three mate pairs due to cryptic relatedness (described earlier) with other mate pairs, resulting in a subset of 21 independent mate pairs included in our assortative mating analysis.

We used an empirical distribution to assess if the observed correlations of ancestry between mate pairs are significantly different from what would be expected under the null hypothesis of random mating. In particular, we randomly permuted the MXL mate pairs 5,000 times, and for each of the 5,000 permutations, we calculated correlations of the mate pairs for each of the three ancestries (European, Native American, and African). The correlations of each ancestry on the autosomes and the X chromosome between mate pairs from the 5,000 permutations were then used to construct empirical distributions under the null hypothesis of random mating in the MXL. The distributions of ancestry correlations among mate pairs are centered around zero when there is random mating, with a standard deviation around 0.2 for each of the three ancestries (Figure S5).

We first tested the null hypothesis versus an alternative hypothesis of assortative mating for ancestry using the observed correlations among mate pairs and the empirical null distributions. Table 4 shows the p-values for the autosomal and X chromosome correlations of African, European and Native American ancestry proportions calculated from the 21 MXL mate pairs. There is significant evidence of assortative mating for European and Native American ancestries on the autosomes in the HapMap MXL, with corresponding p-values of 0.015 and 0.017, respectively. There is also significant evidence for assortative mating based on European and Native American ancestry on X chromosome, with p-values of 0.011 and 0.007, respectively. The p-values remain significant, even after Bonferroni correction for testing three ancestries. There is not significant evidence of assortative mating for African ancestry for either the autosomes or the X chromosomes (p=0.26 and 0.14, respectively). For testing the null hypothesis of random mating versus an alternative hypothesis of non-random, e.g., assortative or dissasortative mating, a two-sided test can be conducted. The p-values for this test are given in Table 4 and are roughly twice the assortative mating p-values. We also performed permutation tests to assess evidence of assortative and non-random mating for 11 HapMap ASW mate pairs with a documented offspring. No significant evidence of assortative mating in the ASW was detected, and ASW p-values for the three ancestries tested are given in Table 4.

**Table 4.** Ancestry Correlation Among Mate Pairs. P-values detecting assortative or disassortative for ancestry among 11 HapMap ASW and 21 HapMap MXL mate pairs, calculated on the autosomes and the X chromosome separately. The p-values are calculated from the empirical distribution created from sampling 5,000 mate pairs at random. Results presented under ‘assortative mating’ tested the hypothesis of no assortative mating, while ‘non-random mating’ tested the hypothesis of neither assortative nor disassortative mating.

|  |  | HapMap ASW |  |  | HapMap MXL |  |  |
| --- | --- | --- | --- | --- | --- | --- | --- |
|  |  | African | European | Native American | African | European | Native American |
| Autosomal | assortative mating | 0.365 | 0.388 | 0.234 | 0.139 | 0.015 | 0.017 |
|  | non-random mating | 0.871 | 0.888 | 0.532 | 0.268 | 0.028 | 0.032 |
| X Chromosome | assortative mating | 0.842 | 0.788 | 0.564 | 0.256 | 0.011 | 0.007 |
|  | non-random mating | 1.000 | 1.000 | 1.000 | 0.530 | 0.024 | 0.013 |

### Ancestry Equilibrium on the X Chromosome Under Random Mating After Initial Admixture Event

We also investigated the number of generations required for males and females to reach ancestry equilibrium on the X chromosome in a randomly mating population. We considered the setting where there is admixing between two ancestral populations and where mate pairs at the initial admixture event consist of males with ancestry entirely from one of the populations and females having ancestry derived from the other population. We then performed a simple computation to estimate proportional ancestry for each generation assuming random mating and after an initial admixing event between founder females and males with the most extreme setting of discordant ancestry between the two sexes at the time of admixture. Figure 6 shows the proportion ancestry by generation in the admixed population for males and females. A recent finding published a similar result, although the initial ancestry proportions considered did not include the extreme proportions as we did here (Goldberg and Rosenberg 2015). We find an equilibrium of 1/2 is reached for autosomal ancestry in males and females in the first generation. Proportional ancestry on the X chromosome for males and females tends to an equilibrium of 2/3 and 1/3 of the founder female and male ancestries, respectively, that is achieved around eight generations after the initial admixing event. This result is not surprising since females contribute 2/3 of the X chromosomes in a population. A recent study developed a model that showed the 2/3 and 1/3 ancestry proportions on the X chromosome in admixed populations derived from two ancestries with a single admixture event may be accurate, but is not correct if the admixing is ongoing (Goldberg and Rosenberg 2015). Nevertheless, whether a single admixture event or ongoing admixture is assumed, the X chromosome and the autosomal chromosomes will not have the same equilibrium ancestry proportions in an admixed population when males and females have different ancestries at the time of the admixture event(s).

**Figure 6.**
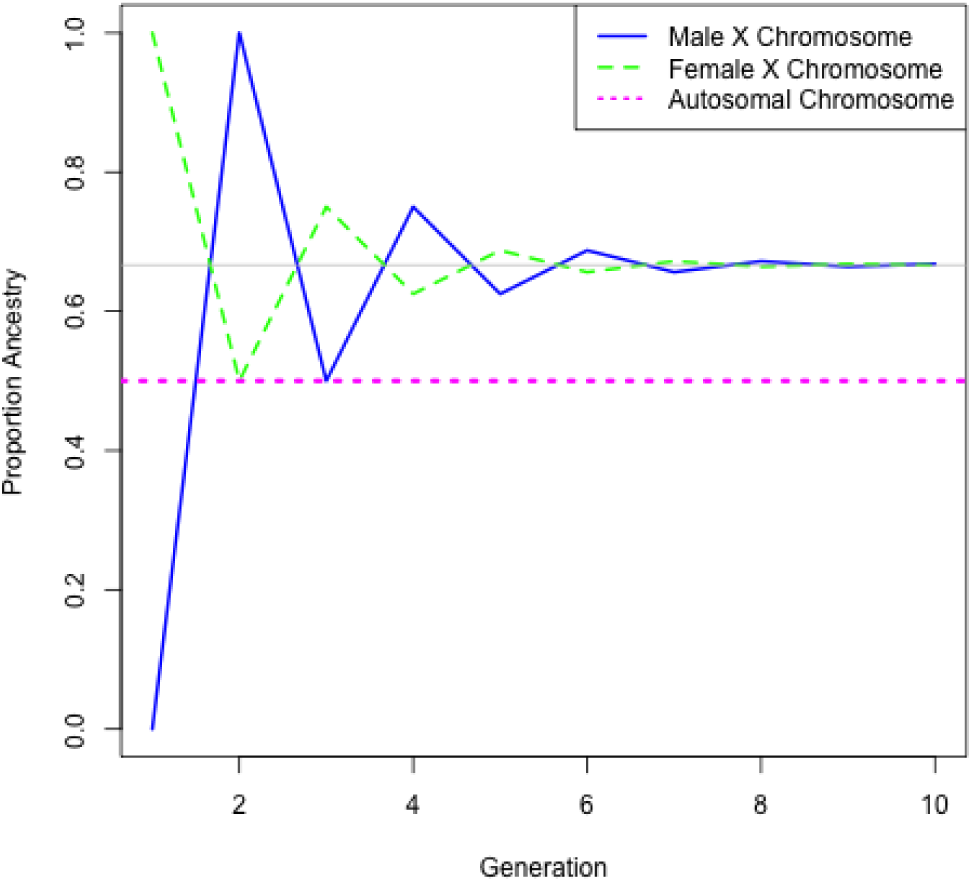
Ancestry Proportions By Generation Under Random Mating. The proportion of ancestry for the autosomes and the X chromosome by sex, assuming females and males have opposite ancestries at the initial admixture event. After the initial admixture event, random mating is assumed. The gray line shows the equilibrium proportions on the X chromosome.

## Discussion

Systematic ancestry differences at genomic loci may arise in recently admixed populations as a result of selection and ancestry related assortative mating. Here, we developed the CAnD method for detecting heterogeneity in population structure across the genome in populations with admixed ancestry. CAnD uses ancestry inferred from SNP genotype data to identify chromosomes that have significantly different contributions from the underlying ancestral populations. The CAnD method takes into account correlated ancestries among chromosomes within individuals for both valid testing and improved power for detecting heterogeneity in population structure across the genome. Some additional features of the CAnD method are: (1) X chromosome data can easily be incorporated in the analysis; and (2) the method can be used for testing heterogeneity in ancestry among any subset of chromosomes in the genome.

We performed simulation studies with admixture and real genotype data from HapMap. We demonstrated that CAnD had appropriate type I error. We also showed in the simulation studies that the CAnD test has higher power to detect heterogeneity in ancestry between chromosomes than a pooled t-test that does not take account correlations in ancestry among chromosomes.

We applied the CAnD method to the HapMap MXL population sample where significant heterogeneity in European ancestry and Native American ancestry was detected across the genome (autosomes and the X chromosome), with p-values of 9e-07 and 1e-06, respectively. A subsequent analysis showed that the heterogeneity in ancestry across the MXL genomes detected by CAnD is largely due to elevated Native American ancestry and deficit of European ancestry on the X chromosomes. These results are consistent with previous reports for U.S. Hispanic/Latinos (Bryc et al. 2015) and Latin Americans (Bryc et al. 2010b), where it has been suggested that the X versus autosomal differences are likely due to sex-specific patterns of gene flow in which European male colonists contributed substantially more genetic material than European females at the time of admixture. There was no significant evidence of genetic heterogeneity in the HapMap ASW detected by CAnD and no significant differences in ancestry between the autosomal chromosomes and the X chromosome were detected. The autosomal chromosomes and the X chromosome in the ASW are largely African derived, and a larger sample is required to have adequate power for the detection of chromosomal ancestry differences in this population.

The CAnD method can incorporate estimates of local ancestry at specific locations across the genomes, using software such as RFMix, or proportional ancestry estimates for each chromosome with software such as FRAPPE or ADMIXTURE. We compared the CAnD results for the HapMap MXL when using local ancestry estimates from RFMix, which requires phased genotype data, to the results when using chromosomal ancestry estimates with FRAPPE where unphased genotype data was used. Heterogeneity in ancestry was detected with CAnD when using either local ancestry estimates from RFMix or chromosomal ancestry estimates from FRAPPE. Interestingly, p-values were slightly smaller when using estimates from FRAPPE that were based on unphased genotype data as compared to using local ancestry estimates from phased genotype data. This result might be an artifact of there being some errors in the phasing and RFMix not appropriately taking into account uncertainty in the phasing when estimating local ancestry.

In the present paper, CAnD was used to identify entire chromosomes with ancestry contributions that are significantly different than expected. If local ancestry estimates are available, CAnD can be used to follow-up on the chromosomal findings by fine-mapping the specific regions that may be under selection. CAnD can be used with a sliding window or a set of genes within a chromosome to localize areas that exhibit heterogeneity in population structure. Future work will consider this approach.

We also investigated the number of generations required for ancestry on the X chromosome to reach equilibrium in males in females after a single admixing event with two populations. In the most extreme setting where all males are from one population and all females are from the other population at the time of admixture, approximately eight generations are required under random mating between males and females to reach ancestry equilibrium on the X. Estimates of the number of generations since admixture in the Mexican population (Johnson et al. 2011) range from 10 to 15. It is reasonable to assume that equilibrium on the X chromosome for males and females should have been reached in the Mexican population if mating in this population is at random. Previous studies (Risch *et al*. 2009; Sebro *et al*. 2010), however, have shown evidence of non-random mating in Mexican populations. In the HapMap MXL, between mate pairs that produced an offspring, we also detected significant evidence of assortative mating, where the correlation of European and Native American ancestries on both the autosomes and the X chromosome is significantly higher than what would be expected under the null hypothesis of random mating. Evaluating differences in ancestry on the X chromosome between males and females may potentially be a useful tool for the detection of non-random mating in recently admixed populations, since under the most extreme setting of discordant ancestry between males and females at the time of admixture, we find that that there should be no difference in ancestry on the X chromosome between males and females after eight generations of random mating. This is future work to be considered.

The CAnD method is implemented in the R language and is available from Bioconductor (http://www.bioconductor.org) as part of the CAnD package.

## Appendix A: Derivation of the Covariance Matrix for CAnD

We initially derive the pairwise covariance of test statistics 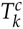, 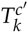 assuming known Σ. Then, we outline how we estimate the parameters required to calculate 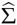 in practice.

Consider individual *i* and subpopulation *k* and denote the ancestry proportion on chromosome *c* as 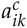. Let *m* be the number of chromosomes under consideration. For chromosomes *c* and *c′*, denote

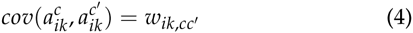

The entries in the 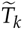. covariance matrix Σ are

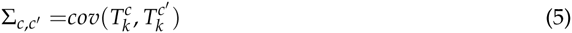

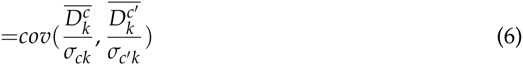

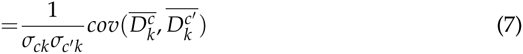

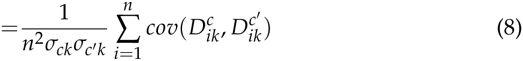

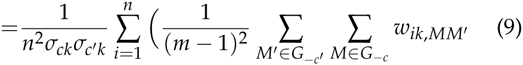

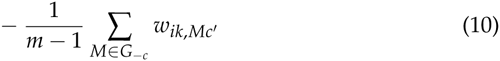

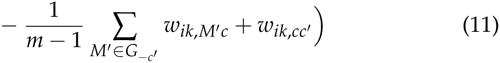

where *G_−c_* is the set of all chromosomes excluding *c*. The diagonal entries correspond to 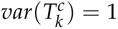.

In practice, we must estimate the values of 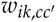 and 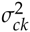. For a given subpopulation *k* and chromosome *c*, denote the average ancestry proportion across all individuals *i* as

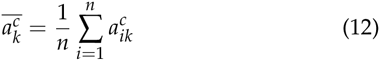

We calculate the covariance of ancestry proportions across individuals in our sample between chromosomes *c* and *c′* as

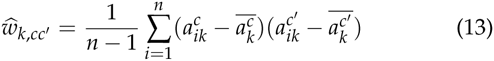

We estimate *σ_ck_*, the standard deviation of 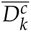, with

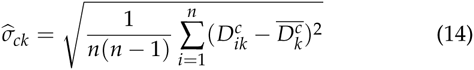

for a given chromosome *c* and subpopulation *k*, and the corresponding element in 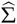 for chromosomes *c* and *c′* is

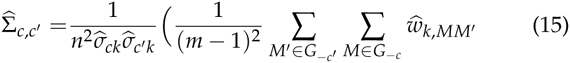

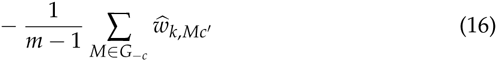

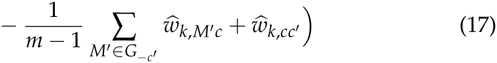

## Literature Cited

Altshuler, D. M., R. A. Gibbs, L. Peltonen, E. Dermitzakis, S. F. Schaffner, et al., 2010 Integrating common and rare genetic variation in diverse human populations. Nature 467: 52–58.

Bhatia, G., A. Tandon, N. Patterson, M. Aldrich, C. B. Ambrosone, et al., 2014 Genome-wide Scan of 29,141 African Americans Finds No Evidence of Directional Selection since Admixture. American Journal of Human Genetics 95: 437–444.

Browning, S. R., and B. L. Browning, 2007 Rapid and accurate haplotype phasing and missing-data inference for whole-genome association studies by use of localized haplotype clustering. American Journal of Human Genetics 81: 1084–1097.

Bryc, K., A. Auton, M. R. Nelson, J. R. Oksenberg, S. L. Hauser, et al., 2010a Genome-wide patterns of population structure and admixture in West Africans and African Americans. Proceedings of the National Academy of Sciences of the United States of America 107: 786–791.

Bryc, K., E. Y. Durand, J. M. Macpherson, D. Reich, and J. L. Mountain, 2015 The genetic ancestry of African Americans, Latinos, and European Americans across the United States. American Journal of Human Genetics 96: 37–53.

Bryc, K., C. Velez, T. Karafet, A. Moreno-Estrada, A. Reynolds, et al., 2010b Genome-wide patterns of population structure and admixture among Hispanic/Latino populations. Proceedings of the National Academy of Sciences 107: 8954–8961.

Goldberg, A. and N. A. Rosenberg, 2015 Beyond 2/3 and 1/3: The Complex Signatures of Sex-Biased Admixture on the X Chromosome. Genetics 201: 263–279.

Jin, W., S. Xu, H. Wang, Y. Yu, Y. Shen, B. Wu, and L. Jin, 2012 Genome-wide detection of natural selection in African Americans pre- and post-admixture. Genome Research 22: 519–527.

Johnson, N. A., M. A. Coram, M. D. Shriver, I. Romieu, G. S. Barsh, et al., 2011 Ancestral components of admixed genomes in a Mexican cohort. PLoS Genetics 7: e1002410.

Li, J. Z., D. M. Absher, H. Tang, A. M. Southwick, A. M. Casto, et al., 2008 Worldwide human relationships inferred from genome-wide patterns of variation. Science 319: 1100–1104.

Manichaikul, A., W. Palmas, C. J. Rodriguez, C. A. Peralta, J. Divers, et al., 2012 Population structure of Hispanics in the United States: The Multi-Ethnic study of Atherosclerosis. PLoS Genetics 8: e1002640.

Maples, B. K., S. Gravel, E. E. Kenny, and C. D. Bustamante, 2013 RFMix: A discriminative modeling approach for rapid and robust local-ancestry inference. American Journal of Human Genetics 93: 278–288.

Nelis, M., T. o. Esko, R. Mägi, F. Zimprich, D. Toncheva, et al., 2009 Genetic structure of Europeans: A view from the NorthEast. PLoS ONE 4: e5472.

Novembre, J., T. Johnson, K. Bryc, Z. Kutalik, A. Boyko, et al., 2008 Genes mirror geography within europe. Nature 456: 98–101.

Price, A. L., A. Tandon, N. Patterson, K. C. Barnes, N. Rafaels, et al., 2009 Sensitive detection of chromosomal segments of distinct ancestry in admixed populations. PLoS Genetics 5: e1000519.

Risch, N., S. Choudhry, M. Via, A. Basu, R. Sebro, et al., 2009 Ancestry-related assortative mating in Latino populations. Genome Biology 10: R132.

Sebro, R., T. J. Hoffman, C. Lange, J. J. Rogus, and N. J. Risch, 2010 Testing for Non-Random Mating: Evidence for Ancestry-Related Assortative Mating in the Framingham Heart Study. Genetic Epidemiology 34: 674–679.

Tang, H., S. Choudhry, R. Mei, M. Morgan, W. Rodriguez-Cintron, et al., 2007 Recent genetic selection in the ancestral admixture of Puerto Ricans. American Journal of Human Genetics 81: 626–633.

Tang, H., J. Peng, P. Wang, and N. J. Risch, 2005 Estimation of individual admixture: Analytical and study design considerations. Genetic Epidemiology 28: 289–301.

Thornton, T., H. Tang, T. J. Hoffmann, H. M. Ochs-Balcom, B. J. Caan, et al., 2012 Estimating kinship in admixed populations. American Journal of Human Genetics 91: 122–138.

Zakharia, F., A. Basu, D. Absher, T. L. Assimes, A. S. Go, et al., 2009 Characterizing the admixed African ancestry of African Americans. Genome Biology 10: R141.

